## Supplementary Figures and Tables for "Rearrangement of 3D genome organization in breast cancer epithelial - mesenchymal transition and metastasis organotropism"

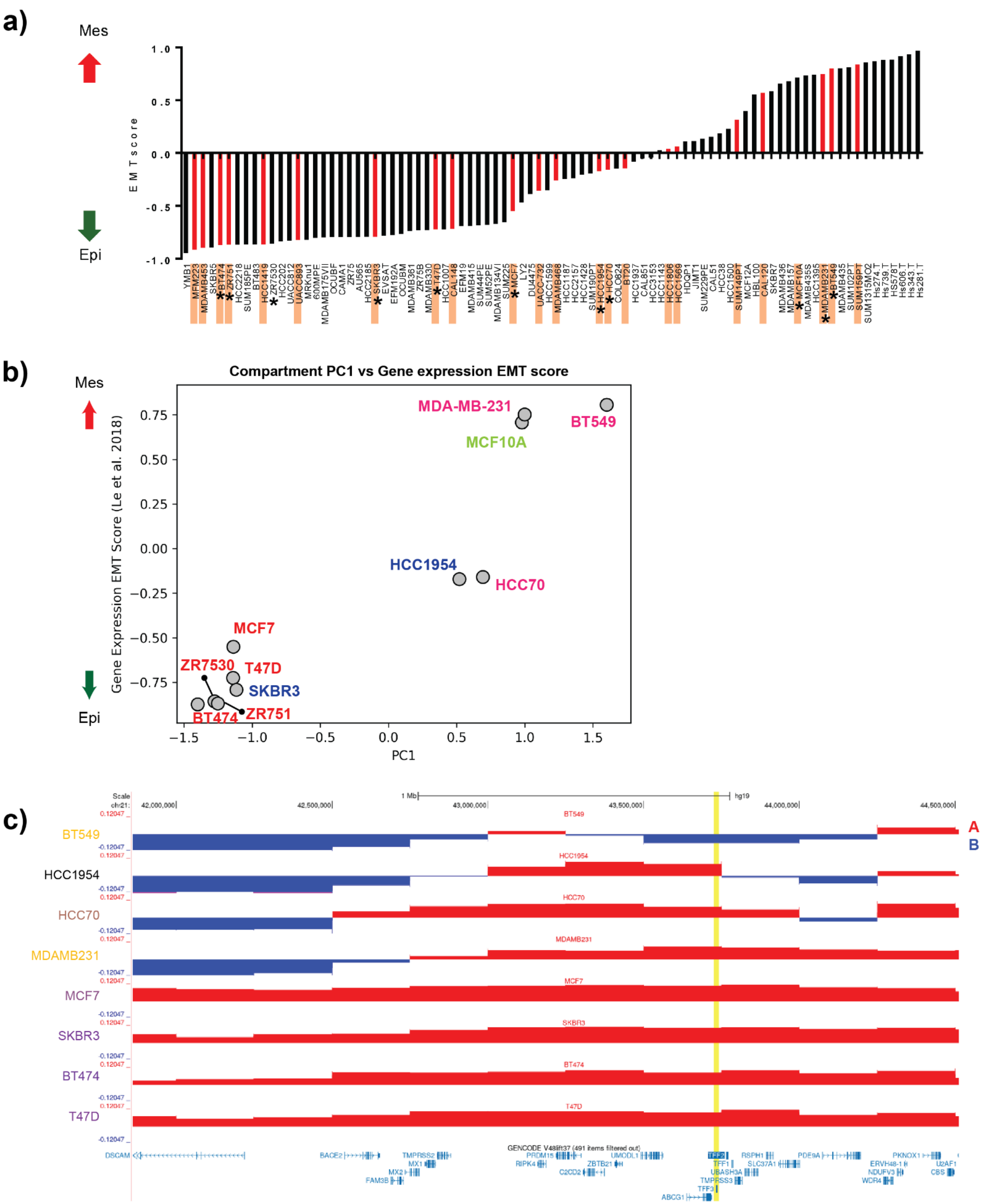


**Supplementary Figure 1. Concordance between EMT classification of cell types by gene expression and Hi-C compartments. a)** Breast cancer cell lines are arranged depending on their epithelial-mesenchymal status. The states are calculated based on their gene expression profile using the method described in Tan et al. (Tan et al., 2014). This figure has been adapted from (Le et al., 2018), reused as consistent with CC-BY License. Stars indicate cell lines used in the current study. **b)** Scatter plot showing relation between compartment PC1 (from Fig. 1c) and gene expression EMT score (as in Supplementary Figure 1a) for breast cancer cell lines considered in this manuscript. **c)** Example of compartment changes near but not on top of a gene that changes expression across the EMT axis. TFF3 (yellow highlighted position) is downregulated in more mesenchymal like cancer cell lines (top 4) as compared to epithelial-like (bottom 4). The gene itself is in the A compartment (red, positive PC1 values) in all but 1 cell line, but many regions nearby switch to the B compartment in the top 4 vs. bottom 4 cell lines.

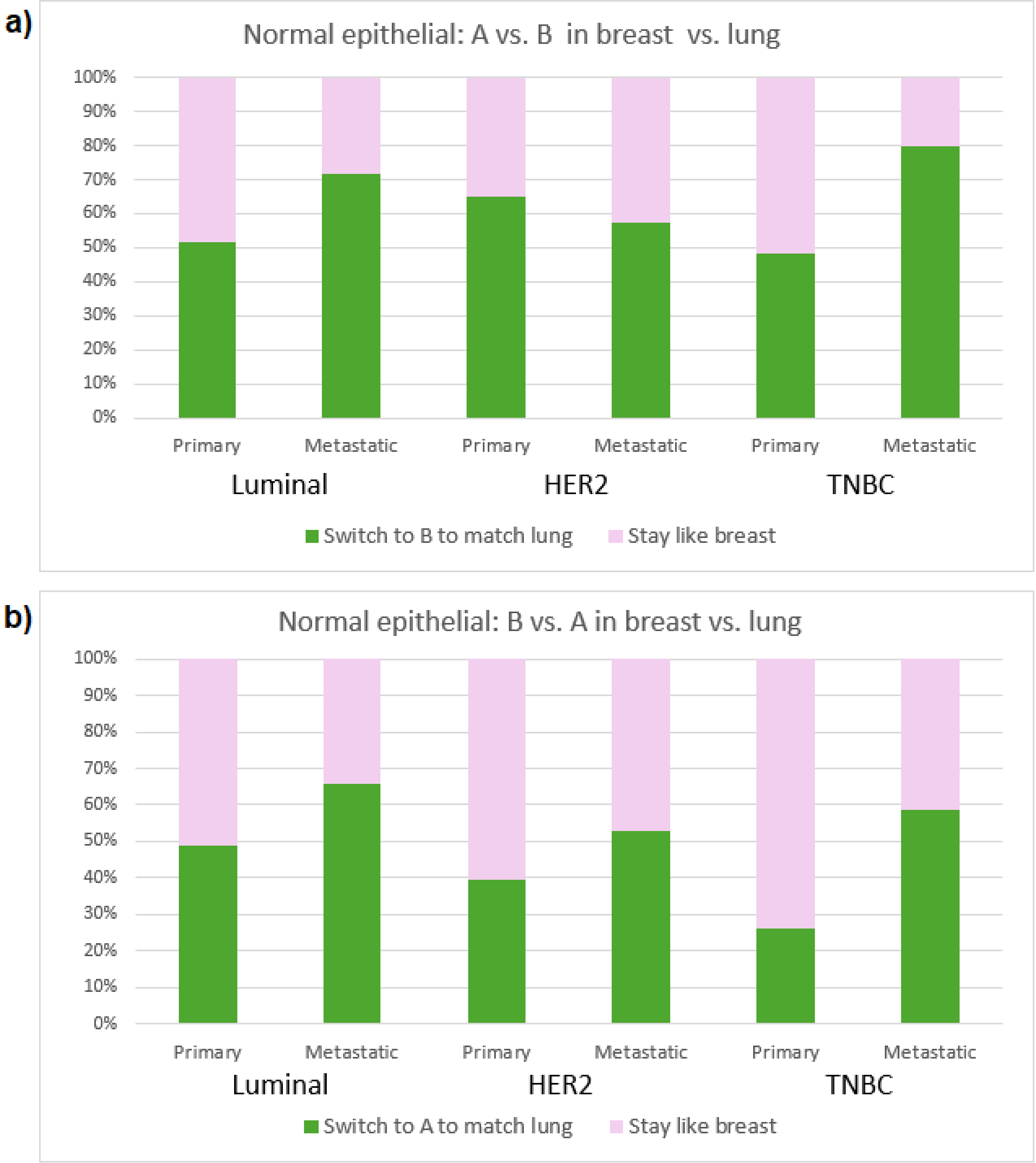


**Supplementary Figure 2. Conservation vs. change in breast cancer cell compartment identity for regions that differ between breast and lung epithelial cells. a)** Out of all genomic bins that are in the A compartment in normal breast epithelial cell lines but B compartment in normal lung epithelial cells, the graph shows the proportion that switch compartments to match lung (green) for each category of breast cancer vs. those that remain like breast (pink). b**)** Same as (a), but for genomic regions that are in the B compartment in normal breast epithelial cells and A compartment in normal lung.

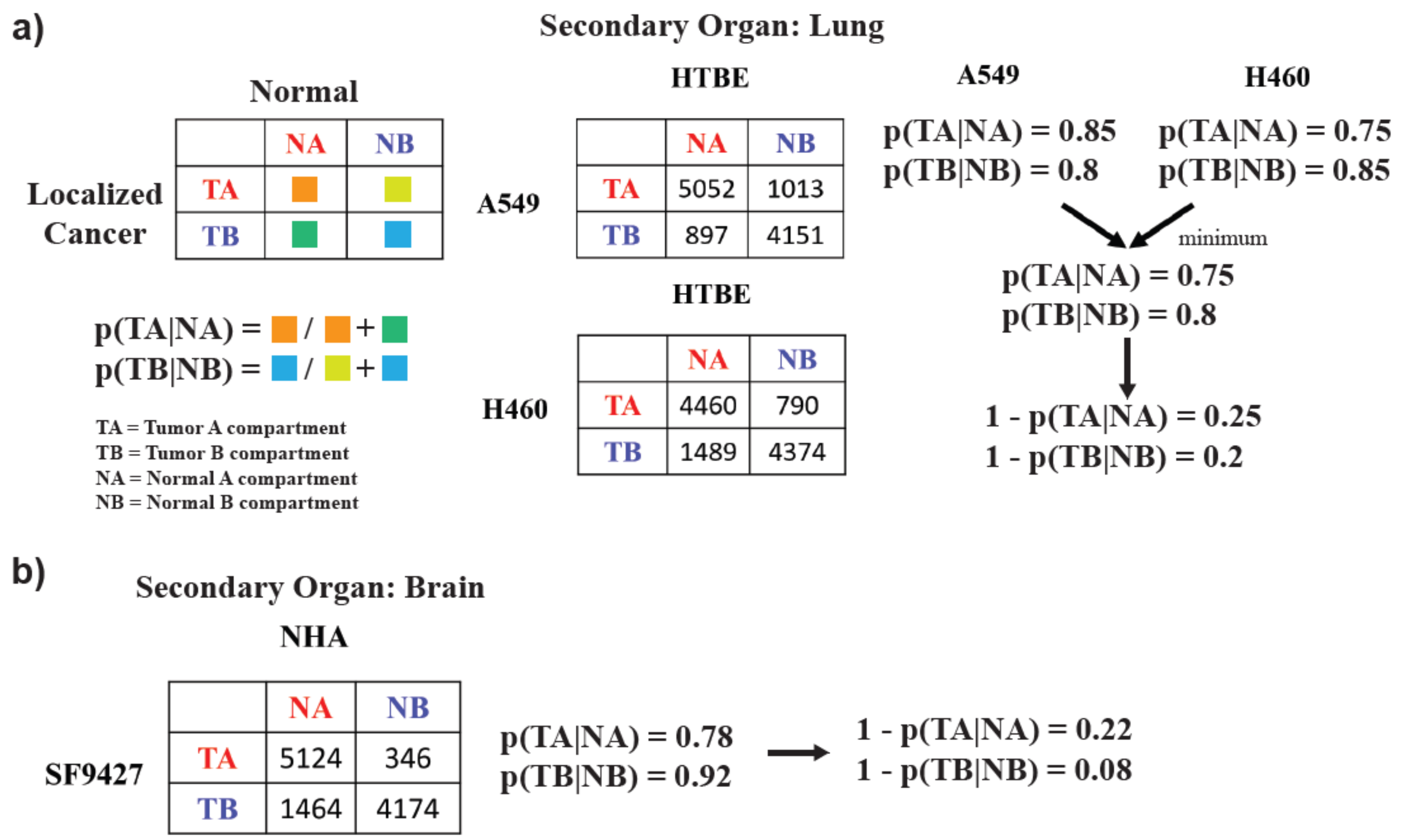


**Supplementary Figure 3. Calculation to adjust organ-permissive compartment switch calculation to account for different background levels of certain compartments. a)** Calculation of the probability of a genomic region to be in a specific compartment in localized lung cancer given that the region belongs to a certain compartment in normal lung epithelial cell. For the A compartment, p(TA|NA) represents the probability of a region in A compartment in localized lung cancer cell given that region also belongs to A compartment in normal lung epithelial cell. Similarly, p(TB|NB) for the B compartment. **b)** Calculation of the probability of a genomic region to be in a specific compartment in localized brain cancer (SF9427) given that the region belongs to a certain compartment in normal brain cell (NHA).

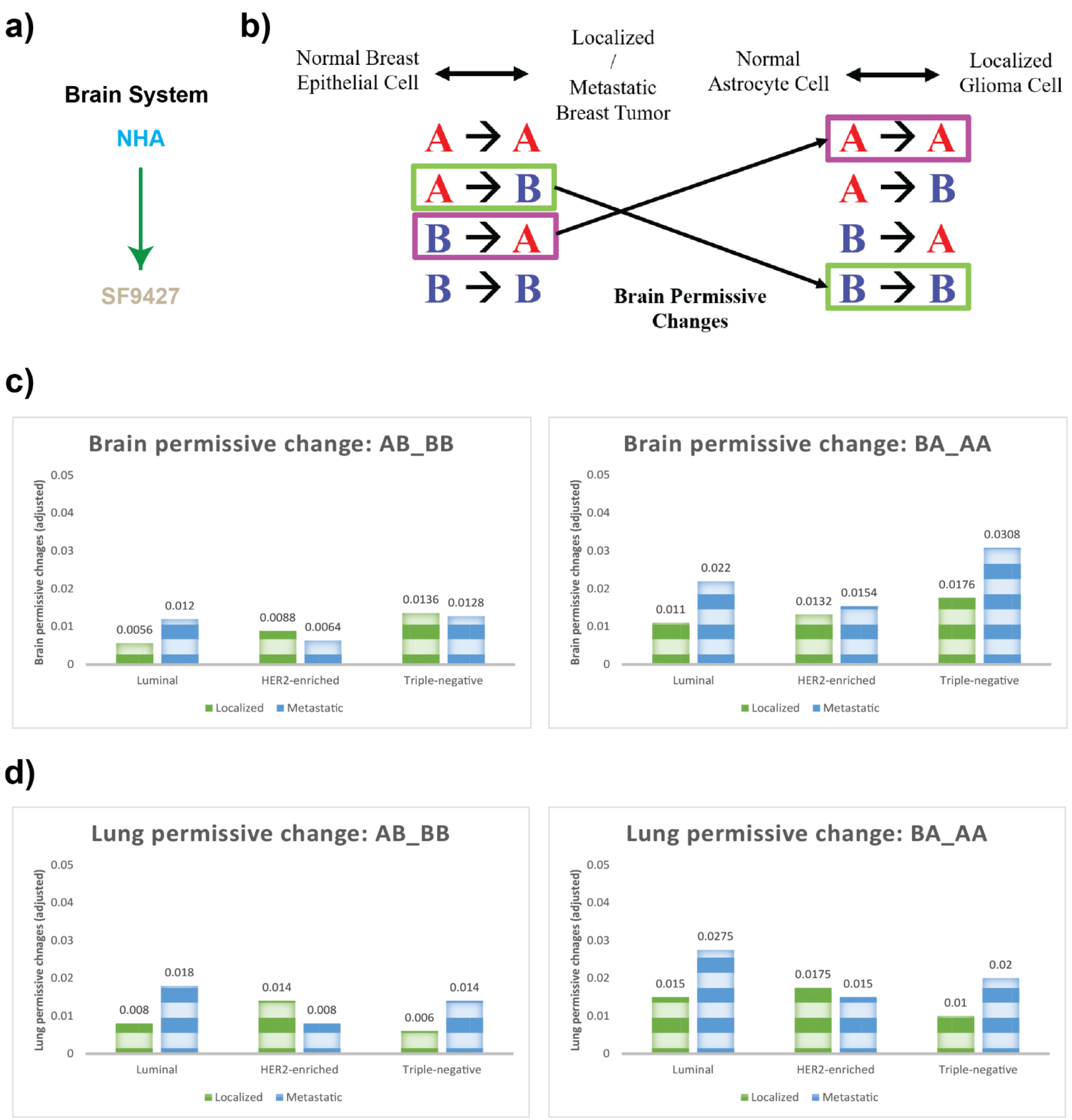


**Supplementary Figure 4. Comparison of Brain-Permissive and Lung-Permissive Changes in Lung Metastatic Breast Cancer. a)** Normal human astrocyte (NHA) and glioblastoma (SF9427) cells are used for compartment comparisons to breast cancer changes. **b)** Schematic diagram representing breast cancer brain-permissive changes calculation. Detailed steps are mentioned in Fig. 4b. **c)** Adjusted (see Supplementary Figure 3 and Methods) levels of different brain permissive changes shown by localized and metastatic cancers from different breast cancer subtypes. **d)** Adjusted levels of different lung permissive changes shown by localized and metastatic cancers from different breast cancer subtypes.

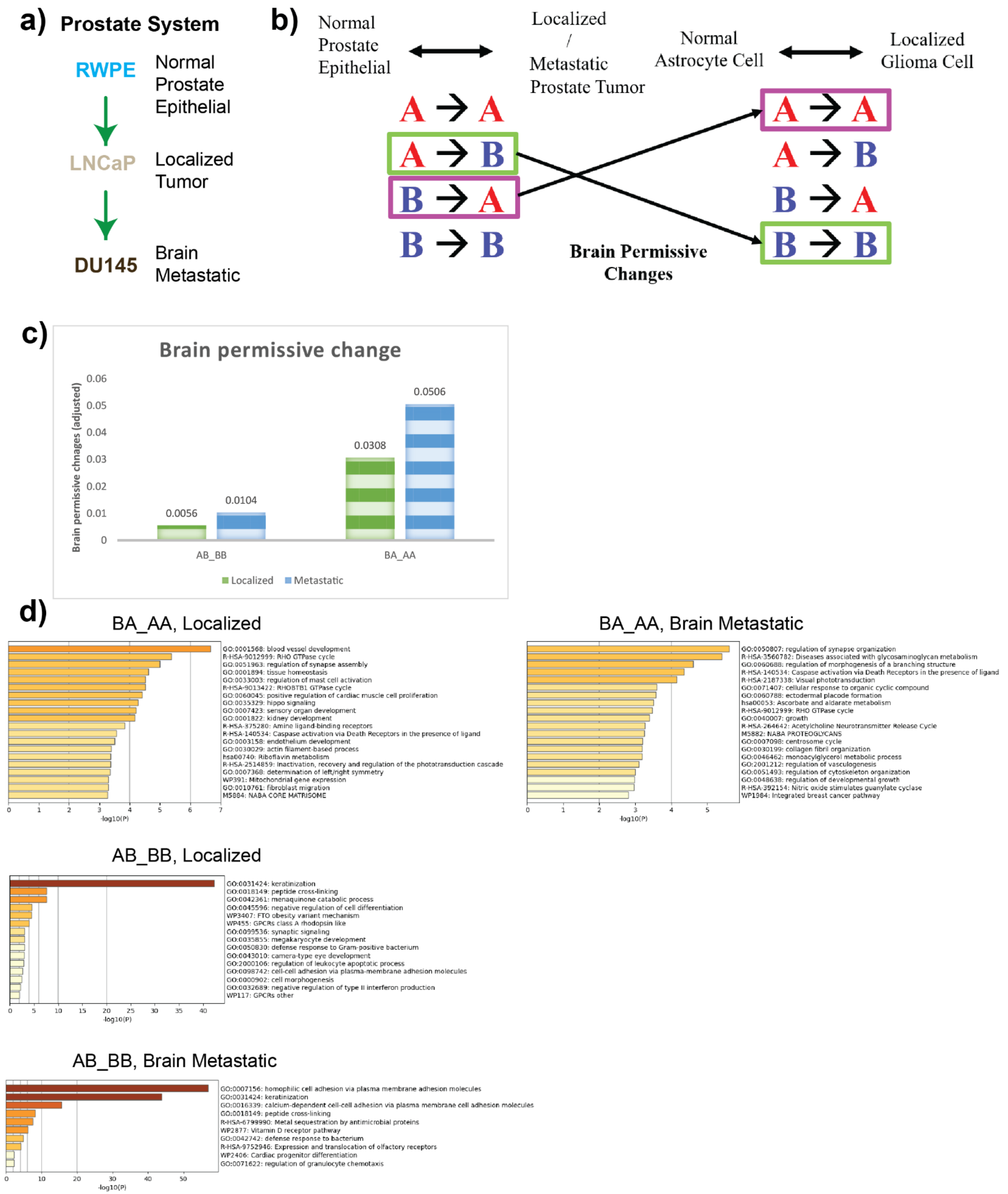


**Supplementary Figure 5. Prostate cancer cells that metastasize to brain show increased brain-permissive compartment changes at neuronal-related genes. a)** Normal epithelial (RWPE), prostate cancer (LNCaP), and brain metastatic prostate cancer (DU145) Hi-C datasets are used for compartment comparisons to glioblastoma/normal brain changes. **b)** Schematic diagram representing prostate cancer brain-permissive changes calculation. Detailed steps are mentioned in Fig. 4b. **c)** Adjusted (see Supplementary Figure 3 and Methods) levels of different brain permissive changes (fraction of compartment changes that are brain permissive) shown by adenocarcinoma-like and brain metastatic prostate cancer. **d)** Gene Ontology term enrichment for genes in regions in each category of comparisons. Note brain-relevant terms such as regulation of synapse organization in genes switched toward the A compartment in cancer that match brain A compartment and keratinization in genes switched toward the B compartment in cancer that match brain B compartment.

| Cell line | Type | Tissue of origin | Race | Collection Site | Subtype | ER | PR | HER2 | Hi-C Accession | Hi-C Replicates | RNA-seq Accession | RNA-seq replicates |
|---|---|---|---|---|---|---|---|---|---|---|---|---|
| HMEC | Human Mammary Epithelium | Breast | Unknown | Breast | Normal | + | + | + | GSE167150 | Replicates Combined | GSE167152 | Replicates 1 and 2 |
| MCF10A | Human Mammary Epithelium | Breast | Caucasian | Breast | Fibrocystic | + | + | + | GSE109229 | Replicates Combined | GSE96860 | Replicates 1, 2, 3 and 4 |
| BT474 | Infiltrative Ductal Carcinoma | Breast | Caucasian | Solid Tumor, breast | Luminal B | + | + | + | GSE109229 | Replicates Combined | GSE179280 | Replicates 1, 2, and 3 |
| ZR751 | Infiltrative Ductal Carcinoma | Breast | Caucasian | Ascites fluid | Luminal A | + | P/N | - | GSE128676 | Replicate 1 | NA | NA |
| ZR7530 | Infiltrative Ductal Carcinoma | Breast | African American | Ascites fluid | Luminal B | + | - | + | GSE167150 | Replicates Combined | GSE167152 | Replicates 1 and 2 |
| MCF7 | Infiltrative Ductal Carcinoma | Breast | Caucasian | Pleural Effusion, Lung | Luminal A | + | + | - | GSE109229 | Replicates Combined | GSE96860 | Replicates 1, 2, 3 and 4 |
| T47D | Infiltrative Ductal Carcinoma | Breast | Caucasian | Pleural Effusion, Lung | Luminal A | + | + | - | GSE167150 | Replicates Combined | GSE167152 | Replicates 1 and 2 |
| HCC1954 | Infiltrative Ductal Carcinoma | Breast | East Indian | Solid Tumor, breast | HER2 | - | - | + | GSE167150 | Replicates Combined | GSE167152 | Replicates 1 and 2 |
| SKBR3 | Adenocarcinoma | Breast | Caucasian | Pleural Effusion, Lung | HER2 | - | - | + | GSE109229 | Replicates Combined | GSE96860 | Replicates 1, 2, 3 and 4 |
| HCC70 | Infiltrative Ductal Carcinoma | Breast | African American | Solid Tumor, breast | Triple Negative A | - | - | - | GSE167150 | Replicates Combined | GSE167152 | Replicates 1 and 2 |
| BT549 | Invasive breast carcinoma | Breast | Caucasian | Solid Tumor, breast | Triple Negative B | - | - | - | GSE167150 | Replicates Combined | GSE167152 | Replicates 1 and 2 |
| MDA-MB-231 | Adenocarcinoma | Breast | Caucasian | Pleural Effusion, Lung | Triple Negative B | - | - | - | GSE143678 | Replicate 1 | GSE179280 | Replicates 1, 2, and 3 |
| HTBE | Bronchial-Tracheal Epithelial cells | Lung | Unknown | Lung | Normal | NA | NA | NA | GSE113703 | Replicates Combined | GSE89008 | Replicates 1 and 2 |
| A549 | Adenocarcinoma | Lung | Caucasian | Solid Tumor, Lung | NSCLC | NA | NA | NA | ENCSR444WCZ | Replicates Combined | ENCSR000CON | Replicates 1 and 2 |
| H460 | Large cell lung cancer | Lung | Unknown | Pleural Effusion, Lung | NSCLC | NA | NA | NA | ENCSR489OCU | Replicates Combined | ENCSR164OCT | Replicates 1 and 2 |
| NHA | Normal Astrocytes | Brain | Unknown | Cerebral cortex | Normal | NA | NA | NA | GSE162976 | Replicate 1 | NA | NA |
| SF9427 | Glioblastoma | Brain | Unknown | Brain | Glioma | NA | NA | NA | GSE162976 | Replicate 1 | NA | NA |
| LNCaP | Adenocarcinoma | Prostate | Caucasian | Lymph Node, Prostate | - | NA | NA | NA | GSM5241744 | Replicate 1 | NA | NA |
| DU145 | Adenocarcinoma | Prostate | Caucasian | Brain | - | NA | NA | NA | GSM5241745, GSM5241746 | Replicates 1 and 2 | NA | NA |

**Supplementary Table 1. Classifications of cell line models used and sources of all publicly available Hi-C and RNA-seq data used in the analyses.** Information about cell lines derived from the following sources: ATCC, Engel and Young 1978, Gazdar et al., 1998, Cellosaurus.org, Wang et al. 2021, and Stone et al., 1978.

**Supplementary Table 2. Lists of genes obtained from compartmental PCA, transcriptomics PCA and curated epithelial-mesenchymal gene set.** Compartmental_PC1_Positive: genes found within the top 100 highly varying genomic regions in compartmental PCA analysis (positive PC1); Compartmental_PC1_Negative: genes found within the top 100 highly varying genomic regions in compartmental PCA analysis (negative PC1); RNAseq_PC1_Positive: top 100 highly varying genes in transcriptomics PCA analysis (positive PC1); RNAseq_PC1_Negative: top 100 highly varying genes in transcriptomics PCA analysis (negative PC1); Curated_Epithelial: curated epithelial genes; Curated_Mesenchymal: curated mesenchymal genes.

| Compartmental PC1_Positive | Compartmental PC1_Negative | RNAseq_PC1 Positive | RNAseq_PC1 Negative | Curated_ Epithelial | Curated_ Mesenchymal |
|---|---|---|---|---|---|
| SERPINB13 | FRMD3 | SCGB2A2 | MT1E | ABCC3 | AKAP12 |
| SERPINB4 | IDNK | TRIL | PRKCDBP | ABHD11 | AKAP2 |
| SERPINB3 | NTN1 | S100A14 | VIM | ADAP1 | AKT3 |
| SERPINB11 | LOC101928266 | IGFBP5 | CAV1 | AGR2 | ANGPTL2 |
| SERPINB7 | STX8 | ALDH3B2 | SNAI2 | CRYBG1 | ANK2 |
| MIRLET7A2 | PEPD | AZGP1 | AXL | AKR1B10 | AP1S2 |
| MIR100 | CHST8 | C1orf64 | MMP14 | ALDH3B2 | ASPN |
| MIR100HG | DNAJC1 | SLC44A4 | GPX1 | ALOX5 | AXL |
| PRNP | EBLN1 | RAB25 | SERPINE1 | ANK3 | BAG2 |
| PRND | TUNAR | KRT4 | PTRF | ANXA4 | BGN |
| PRNT | LINC00551 | INHBB | CXCL1 | ANXA9 | BICC1 |
| ETS1 | LINC00443 | DSCAM-AS1 | EMP3 | AP1M2 | BNC2 |
| MIR6090 | MYH14 | LRRC26 | RGS4 | AQP3 | C1orf54 |
| LOC101929517 | KCNC3 | SPDEF | PNMAL1 | ARAP2 | C1R |
| UBASH3B | NAPSB | CEACAM6 | FOSL1 | AREG | C1S |
| CRTAM | NAPSA | C2orf54 | MSN | ARHGAP32 | CALD1 |
| LOC339593 | NR1H2 | FXYD3 | ZNF655 | ARHGAP8 | CAV1 |
| MIR125B1 | POLD1 | HOXC10 | PLAU | ARHGDIB | CCL2 |
| BLID | SPIB | S100P | SPARC | ARHGEF5 | CCL8 |
| CYYR1 | MYBPC2 | C9orf152 | MSLN | ATP1B1 | CD163 |
| LOC101929413 | FAM71E1 | PLA2G2A | LOX | ATP2C2 | CDH11 |
| SIRPD | EMC10 | FOXA1 | TGFB1I1 | AZGP1 | CDH2 |
| SIRPB1 | SH3GL3 | TMEM125 | FST | B3GNT3 | CDK14 |
| SIRPG | ADAMTSL3 | RHOV | HTRA1 | BCAS1 | CEP170 |
| SIRPG-AS1 | COLEC12 | BCAS1 | RAB34 | BIK | CHN1 |
| PTPRM | CETN1 | LAD1 | GAS6 | BLNK | CHRDL1 |
| MIR5009 | CLUL1 | TFF3 | IGFBP6 | BSPRY | CLEC2B |
| LOC101926942 | TYMSOS | ELF3 | ADAMTS1 | INAVA | CLIC4 |
| KIF20B | TYMS | EFHD1 | AKR1B1 | C1orf116 | COL14A1 |
| LINC00865 | ENOSF1 | SMIM22 | CXCL2 | C4orf19 | COL15A1 |
| LINC01375 | YES1 | CLDN3 | SRGN | CAMK2N1 | COL5A2 |
| CUBN | ADGRE3 | PRR15L | CCDC80 | CAPN1 | COL6A1 |
| TRDMT1 | ZNF333 | CX3CL1 | UCN2 | CBLC | COL6A2 |
| C20orf196 | ADGRE2 | TRPV6 | SERPINE2 | CD2AP | COLEC12 |
| CHGB | OR7C1 | CALML5 | CHST2 | CD46 | CRISPLD2 |
| TRMT6 | OR7A5 | PRSS22 | NT5E | CD9 | CRYAB |
| MCM8 | OR7A10 | IVL | S100A3 | CDH1 | CSF2RB |
| MCM8-AS1 | OR7A17 | ST14 | TGM2 | CDH3 | CSRP2 |
| CRLS1 | INSC | GGT6 | TNFSF9 | CDS1 | CTSK |
| LINC01182 | ACSM2B | TMPRSS13 | NNMT | CEACAM1 | CXCL12 |

| | | | | | |
|---|---|---|---|---|---|
| NALCN | ACSM1 | S100A9 | IFI27L2 | CEACAM5 | CXCL13 |
| ITGBL1 | THUMPD1 | SCGB1D2 | LIX1L | CEACAM6 | CXCR4 |
| HTR7 | LOC102724084 | KCTD15 | APOBEC3C | CEACAM7 | CYP1B1 |
| RPP30 | DYNLRB2 | TMEM30B | COL6A1 | CLDN3 | DCN |
| ANKRD1 | LINC01227 | CRB3 | GNG11 | CLDN4 | DDR2 |
| XLOC_008559 | CDYL2 | CLDN8 | ALDH3A1 | CLDN7 | DENND5A |
| MTCL1 | GPR26 | ENTPD2 | TNFRSF10D | CNKSR1 | DPT |
| CCBE1 | CPXM2 | PNMT | ACSL4 | COMT | DPYSL3 |
| NRG1-IT3 | LOC102724710 | KLHDC7A | IFI16 | CORO2A | DSE |
| NRG1 | FLJ46284 | KCNJ11 | WBP5 | CTSH | ECM2 |
| LINC01048 | ZMAT4 | AGR2 | S100A2 | CXADR | EFEMP1 |
| LINC00547 | MSX2 | HOXC13 | TPM2 | CYB561 | EFEMP2 |
| POSTN | MIR4634 | EPHB3 | FOXQ1 | CYP4F3 | EMP3 |
| TRPC4 | LOC102724957 | HID1 | COL4A1 | DDR1 | ENPP2 |
| ADAMTS1 | LINC00470 | HOXC11 | LOXL2 | DENND2D | ETV1 |
| LOC100289473 | MIPOL1 | ENPP5 | PTX3 | DHCR24 | EVI2A |
| SIRPA | LOC101927847 | ID4 | FOXC1 | DSC2 | F13A1 |
| PDYN | FOXA1 | DEGS2 | ETS1 | DSG2 | FAP |
| MIR5580 | TTC6 | DLX3 | IGFBP7 | DSP | FBLN1 |
| BMP4 | PCAT18 | PRLR | KIRREL | DTX4 | FBN1 |
| LOC101929095 | AQP4 | CCDC64B | SH3RF3-AS1 | EHF | FERMT2 |
| C1QTNF7 | AQP4-AS1 | MCF2L-AS1 | CA9 | ELF3 | FGL2 |
| CC2D2A | CHST9 | MAL2 | MAP7D3 | ELF5 | FHL1 |
| APP | VRK3 | EPB41L4A-AS2 | MRC2 | ELMO3 | FLI1 |
| IL21 | ZNF473 | PRSS8 | MYL9 | EPB41L4B | FLRT2 |
| IL21-AS1 | FLJ26850 | C1orf210 | SDPR | EPCAM | FN1 |
| CETN4P | SNAR-A8 | PRODH | TIMP1 | EPHA1 | FSTL1 |
| BBS12 | SNAR-A3 | TINCR | FN1 | EPN3 | FXYD6 |
| FGF2 | SNAR-A5 | SPTSSB | NOG | EPS8L1 | FYN |
| SORL1 | SNAR-A14 | AP1M2 | PLAT | EPS8L2 | GAS1 |
| LINC00502 | SNAR-A4 | EPN3 | CCDC8 | ERBB2 | GEM |
| NUDT9P1 | SNAR-A6 | PVRL4 | PNMA2 | ERBB3 | GFPT2 |
| PCGF5 | SNAR-A7 | KLHL9 | CXCL3 | ERMP1 | GIMAP4 |
| ST3GAL6 | SNAR-A9 | BSPRY | SH2D5 | ESRP1 | GIMAP6 |
| DCBLD2 | SNAR-A11 | SYNE4 | LSP1 | ESRP2 | GJA1 |
| LINC01085 | SNAR-A10 | SLC40A1 | COL6A2 | EVPL | GLIPR1 |
| PAPPA | SNAR-B1 | MZB1 | EVI2A | EXPH5 | GLYR1 |
| CAMK4 | SNAR-B2 | YBX2 | MARVELD1 | EZR | GNG11 |
| LOC101928775 | SNAR-D | TC2N | TGFB1 | F11R | GPM6B |
| LINC00474 | IZUMO2 | ATP6V1B1 | TGFBI | FA2H | GREM1 |
| TNC | MIR184 | GPX2 | AHNAK2 | FAM174B | GSC |
| LOC101928748 | ANKRD34C | PRR15 | PROCR | FBP1 | GUCY1B1 |
| DEC1 | ANKRD34C-AS1 | RBBP8NL | IRX1 | FGFR3 | GZMK |
| LINC01080 | TMED3 | ESRP1 | CAV2 | FOXA1 | HEG1 |
| MIR548G | KIAA1024 | SLC52A3 | B3GNT5 | FOXC2 | IFFO1 |
| COL8A1 | CTD-2151A2.1 | HIST1H2BG | GLIPR1 | FUT2 | IGF1 |
| TSLP | TCAF2P1 | SLC2A10 | TMEM158 | FUT3 | IGFBP5 |
| WDR36 | LOC154761 | HIST1H2AE | ITGA5 | FXYD3 | IL10RA |

| | | | | | |
|---|---|---|---|---|---|
| SMOX | TCAF1 | CRISP3 | PTGS2 | GALE | ISLR |
| LINC01433 | OR2F2 | ALDH2 | KRT14 | GALNT3 | ITM2A |
| ADRA1D | OR2F1 | OVOL1 | C3 | GALNT7 | JAM2 |
| SLX4IP | OR6B1 | ATP2A3 | FNDC4 | GDF15 | JAM3 |
| JAG1 | OR2A5 | PADI2 | LGALS1 | GMDS | KCNJ8 |
| MIR6870 | TFF2 | BPIFB1 | LINC00941 | ADGRG1 | POGLUT2 |
| LOC101929395 | TFF1 | FMOD | HLA-A | GPRC5A | JCAD |
| LINC00911 | TMPRSS3 | TACSTD2 | PHLDA1 | GPX2 | KLF8 |
| FLRT2 | UBASH3A | INPP5J | FLRT2 | GRB7 | P3H1 |
| LOC100192426 | RSPH1 | C8orf4 | ANGPTL4 | GRHL1 | LGALS1 |
| HAS2 | SLC37A1 | MARVELD3 | MXRA7 | GRHL2 | LHFPL6 |
| HAS2-AS1 | SEMA4D | NCCRP1 | GAS6-AS2 | GRHL3 | LOX |
| TMEM230 | GADD45G | | | HDHD3 | LOXL2 |
| PCNA | PRCAT47 | | | HES1 | LY96 |
| PCNA-AS1 | C8orf4 | | | HNMT | MAF |
| CDS2 | OLIG2 | | | HPGD | MAFB |
| PAK3 | LINC00945 | | | ICA1 | MAP1B |
| CAPN6 | OLIG1 | | | IL1RN | MEOX2 |
| SPDL1 | LOC101928107 | | | IL20RA | MFAP4 |
| DOCK2 | TOX3 | | | IRF6 | MMP2 |
| LOC101929217 | CASC16 | | | ITGB4 | MOXD1 |
| LOC101929268 | SEC23B | | | ITGB6 | MPDZ |
| EFCAB1 | LINC00493 | | | JUP | MS4A4A |
| LOC100507534 | DTD1 | | | KCNK1 | MS4A6A |
| LOC101927132 | LOC101929526 | | | KLF5 | MSN |
| ADAMTS5 | LSM14A | | | KLK6 | MXRA7 |
| MIR4759 | KIAA0355 | | | KRT15 | MYH10 |
| STARD4 | GGT1 | | | KRT18 | MYL9 |
| STARD4-AS1 | BCRP3 | | | KRT19 | MYLK |
| FAS | POM121L10P | | | KRT7 | NAP1L3 |
| FAS-AS1 | PIWIL3 | | | KRT8 | NR3C1 |
| MIR4679-2 | TOP1P2 | | | LAD1 | NUAK1 |
| MIR4679-1 | SGSM1 | | | LCN2 | OLFML2B |
| CH25H | LOC101927843 | | | LLGL2 | OLFML3 |
| LIPA | LINC00308 | | | LRRC1 | PCOLCE |
| ACTA2 | SYK | | | LSR | PDGFC |
| MIR4666B | NUDT7 | | | LY75 | PDZRN3 |
| IL1RAPL1 | VAT1L | | | MALL | PLEKHO1 |
| ANXA5 | RGS6 | | | MANSC1 | PLN |
| TMEM155 | MIR7843 | | | MAP7 | PLXNC1 |
| PP12613 | RORA | | | MAPK13 | PMP22 |
| EXOSC9 | PCAT19 | | | MGST2 | POPDC3 |
| CCNA2 | LINC01480 | | | MLPH | PTGDS |
| BBS7 | CEACAM21 | | | MPZL2 | PTGIS |
| ATG3 | CEACAM4 | | | MST1R | PTPRC |
| SLC35A5 | CEACAM7 | | | MTUS1 | CAVIN1 |
| LINC01279 | CEACAM5 | | | MUC1 | PTX3 |
| CCDC80 | VPS13C | | | MYH14 | QKI |
| LOC101929694 | GAREM | | | MYO1D | RARRES2 |

| | | | | | |
|---|---|---|---|---|---|
| LOC101928437 | WBP11P1 | | | MYO5C | RECK |
| LINC01023 | PGR | | | MYO6 | RGS2 |
| FER | LOC101054525 | | | NQO1 | RUNX1T1 |
| CASC17 | LINC00652 | | | OAS1 | SACS |
| LAMA2 | C20orf78 | | | OCLN | SAMSN1 |
| ARHGAP18 | SCP2D1 | | | OR7E14P | SDC2 |
| LINC00504 | LOC100270804 | | | OVOL1 | SERPINE1 |
| LINC01038 | LOC400655 | | | OVOL2 | SERPINF1 |
| LINC00382 | LOC100505817 | | | PERP | SERPING1 |
| ZC3H12C | FBXO15 | | | PHLDA2 | SFRP1 |
| CYSLTR2 | LDLRAD4 | | | PKP3 | SFRP4 |
| LOC102723505 | LDLRAD4-AS1 | | | PLLP | SH2B3 |
| LINC01152 | MIR5190 | | | PLS1 | SLC2A3 |
| LOC102723517 | OR7C2 | | | PLXNB2 | SLIT2 |
| SOX9-AS1 | SLC1A6 | | | POF1B | SNAI1 |
| LOC101928205 | CCDC105 | | | PLPP2 | SNAI2 |
| SOX9 | CASP14 | | | PPFIBP2 | SOAT1 |
| ANK2 | OR1I1 | | | PPL | SOBP |
| CAMK2D | SYDE1 | | | PRR15L | SPARC |
| ALG13 | ILVBL | | | PRSS8 | SPARCL1 |
| TRPC5 | UNQ6494 | | | PSCA | SPOCK1 |
| TRPC5OS | SNHG10 | | | PTK6 | SRGN |
| | GLRX5 | | | PTPRF | SRPX |
| | TCL6 | | | PYCARD | STON1 |
| | TCL1B | | | RAB11FIP1 | SYNE1 |
| | TCL1A | | | RAB20 | SYNM |
| | DSCAM-IT1 | | | RAB25 | SYT11 |
| | DSCAM | | | RABGAP1L | TAGLN |
| | NAA20 | | | RAPGEFL1 | TCF4 |
| | CRNKL1 | | | RBM47 | TGFB1I1 |
| | CFAP61 | | | RHOD | TMEFF1 |
| | MIR8084 | | | RNF128 | TMEM158 |
| | C8orf87 | | | S100A14 | TNC |
| | MX2 | | | S100P | TNS1 |
| | MX1 | | | SCEL | TPM2 |
| | TMPRSS2 | | | SCNN1A | TRPC1 |
| | SLC24A3 | | | SDC1 | TUBA1A |
| | NETO1 | | | SDC4 | TUBB6 |
| | KCTD15 | | | SERINC5 | TWIST1 |
| | LOC286370 | | | SFN | TWIST2 |
| | MIR4290 | | | SH2D3A | UCHL1 |
| | LOC101929374 | | | SH3YL1 | VCAM1 |
| | LOC388882 | | | SHANK2 | VCAN |
| | IGLL1 | | | SLC16A5 | VIM |
| | DRICH1 | | | SLC22A18 | VSIG4 |
| | GUSBP11 | | | SLC35A3 | WIPF1 |
| | LOC100130264 | | | SLC37A1 | WWTR1 |
| | LINC00323 | | | SLC44A4 | ZCCHC24 |
| | MIR3197 | | | SLC9A3R1 | ZEB1 |

| | | | | | |
|---|---|---|---|---|---|
| | BACE2 | | | SLPI | ZEB2 |
| | PLAC4 | | | SORD | ZFPM2 |
| | FAM3B | | | SORL1 | FAM216A |
| | SOD1 | | | SPAG1 | SYNE3 |
| | SCAF4 | | | SPDEF | MRC1 |
| | HUNK | | | SPINT1 | TCF3 |
| | THOC1 | | | SPINT2 | |
| | DSCAM-AS1 | | | SSH3 | |
| | LINC01501 | | | ST14 | |
| | DIRAS2 | | | STAP2 | |
| | LINC01508 | | | STYK1 | |
| | MIR4760 | | | SYNGR2 | |
| | MIS18A | | | TACSTD2 | |
| | MRAP | | | TFF1 | |
| | URB1 | | | TFF3 | |
| | SNORA80A | | | TGFA | |
| | WRB | | | TJP2 | |
| | LCA5L | | | TJP3 | |
| | SH3BGR | | | TMC5 | |
| | MIR6508 | | | TMC6 | |
| | B3GALT5-AS1 | | | TMEM30B | |
| | B3GALT5 | | | TMPRSS2 | |
| | PCP4 | | | TMPRSS4 | |
| | LINC00159 | | | TNFSF13 | |
| | IGSF5 | | | TOB1 | |
| | | | | TOM1L1 | |
| | | | | TOX3 | |
| | | | | TPD52 | |
| | | | | TRIM31 | |
| | | | | TRPM4 | |
| | | | | TSPAN1 | |
| | | | | TSPAN13 | |
| | | | | TSPAN15 | |
| | | | | TSPAN8 | |
| | | | | TTC39A | |
| | | | | TUFT1 | |
| | | | | UGT1A1 | |
| | | | | VAMP8 | |
| | | | | VAV3 | |
| | | | | VGLL1 | |
| | | | | XBP1 | |
| | | | | ZNF165 | |
| | | | | ADIRF | |
| | | | | MISP | |
| | | | | CD24 | |
| | | | | CKMT1A | |

**Supplementary Table 3. Quantifying overlap between EMT and organotropic compartment switched genomic regions.** Each entry represents the fraction of the top 100 positive (left) or negative (right) compartment PC1 regions which are also found in the set of genomic bins that show organotropic compartment switches in the categories shown.

| | PC1 Positive 100 Significant regions | PC1 Negative 100 Significant regions |
|---|---|---|
| Luminal Primary AB_BB + BA_AA | 0.03 | 0.03 |
| Luminal Metastatic AB_BB + BA_AA | 0.03 | 0.03 |
| Her2 Primary AB_BB + BA_AA | 0.03 | 0.02 |
| Her2 Metastatic AB_BB + BA_AA | 0.03 | 0.02 |
| TNBC Primary AB_BB + BA_AA | 0 | 0 |
| TNBC Metastatic AB_BB + BA_AA | 0.03 | 0 |
